# Mediator subunit Med15 dictates the conserved “fuzzy” binding mechanism of yeast transcription activators Gal4 and Gcn4

**DOI:** 10.1101/840348

**Authors:** Lisa M. Tuttle, Derek Pacheco, Linda Warfield, Steven Hahn, Rachel E. Klevit

**Affiliations:** Division of Basic Sciences, Fred Hutchinson Cancer Research Center, Seattle, WA 98109, USA; Department of Biochemistry, University of Washington, Seattle, WA 98195, USA

**Keywords:** transcription activation, intrinsically disordered proteins, fuzzy binding

## Abstract

The acidic activation domain (AD) of yeast transcription factor Gal4 plays a dual role in both transcription repression and activation through sequence-dependent binding to Gal80 repressor and sequence-independent binding to Mediator subunit Med15. The activation function of Gal4 arises from two hydrophobic regions within the 40-residue AD. We show by NMR that each AD region binds the Mediator subunit Med15 using a “fuzzy” protein interface. Remarkably, comparison of chemical shift perturbations shows that Gal4 and Gcn4, two ADs of different sequence, interact nearly identically with Med15. The findings that two ADs of different sequence use an identical fuzzy binding mechanism shows a common sequence-independent mechanism for AD-Mediator binding, similar to interactions within a hydrophobic cloud. In contrast, the same region of Gal4 AD interacts with Gal80 via a tight structured complex, implying that the structured binding partner of an intrinsically disordered protein dictates the type of protein interaction.

## INTRODUCTION

The regulation of yeast Gal4-dependent genes is a classic model system for the study of eukaryotic gene activation (Hahn and Young, 2011; Ptashne and Gann, 2002). In the absence of galactose, Gal4 activation function is blocked by the repressor Gal80, while in the presence of galactose, Gal4 activates the transcription of genes required for galactose catabolism. Gal4 consists of an N-terminal DNA binding domain that targets the upstream activating sequence (UAS) of Gal4-regulated genes, a large central domain that likely plays a regulatory role, and a C-terminal transcription activation domain (AD) (Keegan et al., 1986; Ma and Ptashne, 1987b; Sood and Brickner, 2017; Stone and Sadowski, 1993; Wu et al., 1996). Under non-inducing conditions, Gal80 binds the Gal4 AD tightly and blocks its function (Johnston et al., 1987; Ma and Ptashne, 1987a). Upon binding galactose and ATP, the transducer protein Gal3 binds Gal80 to inhibit repressor function, permitting the Gal4 AD to activate transcription (Lavy et al., 2012; Li et al., 2010; Sil et al., 1999). Gal4 was the first transcription factor shown to function in yeast, plants, insects, and animals, demonstrating wide conservation of activation mechanisms across eukaryotes (Fischer et al., 1988; Kakidani and Ptashne, 1988; Ma et al., 1988; Webster et al., 1988).

The Gal4 AD, residues 840-881, is conserved among yeasts, is predicted to be intrinsically disordered, and contains several short hydrophobic stretches set in a background of acidic residues (**Fig. S1**). Residues 855-870 are the most highly conserved and play direct roles in both repression, through binding to Gal80 repressor, and activation. Despite this dual function, mutagenesis studies of Gal4 have revealed a fundamental difference in the requirements for repression and transcription activation. While individual mutations in five Gal4 residues within the AD block Gal80 repression, no single residue is critical for its activation function as mutation of nearly every residue within the AD is tolerated (Ansari et al., 1998; Salmeron et al., 1990). Furthermore, the Gal4-Gal80 interaction is high affinity (K_d_ in the nM range; (Wu et al., 1996)) and the Gal4 AD peptide binds as an α-helix within a cleft formed by the N and C-terminal Gal80 domains (Kumar et al., 2008; Thoden et al., 2008). Altogether, the existing information suggests that Gal4-Gal80 binding involves a sequence-specific interface, while AD function does not.

The Med15 subunit of the transcription coactivator Mediator is important for activation of Gal4 target genes and binding of Gal4 to Mediator requires the Med15 subunit (Jeong et al., 2001; Suzuki et al., 1988). Crosslinking studies showed that the Gal4 AD interacts with Med15 during transcription activation and biochemical analysis identified the N-terminus of Med15 as the Gal4-binding region (Jeong et al., 2001; Reeves and Hahn, 2005). How Gal4-Med15 binding occurs in the apparent absence of sequence-specific interactions and what properties allow the same region of Gal4 to bind different proteins through fundamentally different binding modes remain unanswered questions.

Acidic ADs are found in a large class of transcription factors, many of which regulate inducible genes (Erkina and Erkine, 2016; Hahn and Young, 2011). Acidic ADs are thought to be intrinsically disordered, have functionally important hydrophobic residues embedded within regions of net negative charge, and often target multiple unrelated subunits of transcription coactivators such as Mediator, SAGA, Swi/Snf, etc. The best studied activator of this class is yeast Gcn4, which contains tandem ADs that, like Gal4, bind to Med15 (Fishburn et al., 2005; Herbig et al., 2010; Jackson et al., 1996; Jedidi et al., 2010). The activating function of the Gcn4 C-terminal AD is largely contained within a short 5-residue-long sequence embedded within an acidic background, while the N-terminal AD consists of 4 hydrophobic segments, distributed over 100 residues within a disordered acidic polypeptide (Brzovic et al., 2011; Jackson et al., 1996; Staller et al., 2018; Tuttle et al., 2018; Warfield et al., 2014). Each Gcn4 AD can bind any of several activator-binding domains (ABDs) within Med15 via a fuzzy interface, in which interactions are highly dynamic and no unique protein-protein interface exists. In the large complex of tandem Gcn4 ADs and the complete activator-binding region of Med15, fuzzy binding is retained with nearly every possible AD-ABD interaction detected in a mechanism termed a “fuzzy free-for-all” (Tuttle et al., 2018). Whether the fuzzy-free-for-all mechanism is employed more generally by other transcription factors is an important open question.

We have investigated the interactions between Gal4 AD and Med15 using a combination of biochemical, molecular genetics, and NMR analysis and find that Gal4 AD-Med15 interactions involve a dynamic fuzzy binding mechanism. Despite the many sequence and mechanistic differences involved in Gal4 regulation, we find remarkable similarity in the interfaces of the Med15 ABDs with Gcn4 and Gal4, implying conservation of a sequence-independent interaction that involves a “cloud” of hydrophobic character. By extrapolation, our results suggest that a fuzzy binding mechanism is conserved among many acidic activators and provide a rationale for how multiple transcription activators with different primary sequences can target the same set of activator-binding domains.

## RESULTS

### Multiple hydrophobic regions of the Gal4 AD are important for transcription activation

The Gal4 AD (C-terminal residues 840-881) contains short hydrophobic segments embedded within an acidic background: defined as Region 1 (residues 840 – 851) and Region 2 (residues 855-869), where each contains at least three aromatic residues (Trp, Tyr, or Phe) (Fig. 1; **Fig. S1**). A second segment with AD potential (Gal4 residues 148-196) is situated near the DNA binding domain but does not appear to function in the context of full-length Gal4 (Ma and Ptashne, 1987b). As there is some controversy regarding which residues contribute to Gal4 AD function (Piskacek et al., 2017), we first identified residues required for activation in vivo. We used a model system, in which Gal4 residues 828-881 were fused to the N-terminal linker region and DNA-binding domain of Gcn4 (residues 132-281). This Gcn4 derivative has no inherent AD function and can accept a wide variety of natural and synthetic ADs to allow activation of yeast Gcn4-dependent genes (Pacheco et al., 2018; Warfield et al., 2014). We used a Gal4 AD segment longer than the minimal AD because initial studies indicated that residues 840-881 formed a hydrogel upon purification while residues 828-881 remain soluble at very high concentrations, permitting structural, biochemical and molecular analyses using the same Gal4 AD residues. Gal4 AD function was measured at the Gcn4-dependent genes *ARG3* and *HIS4* after addition of sulfometuron methyl (SM) to induce amino acid starvation and induction of Gal4-Gcn4 protein synthesis.

**Figure 1.**
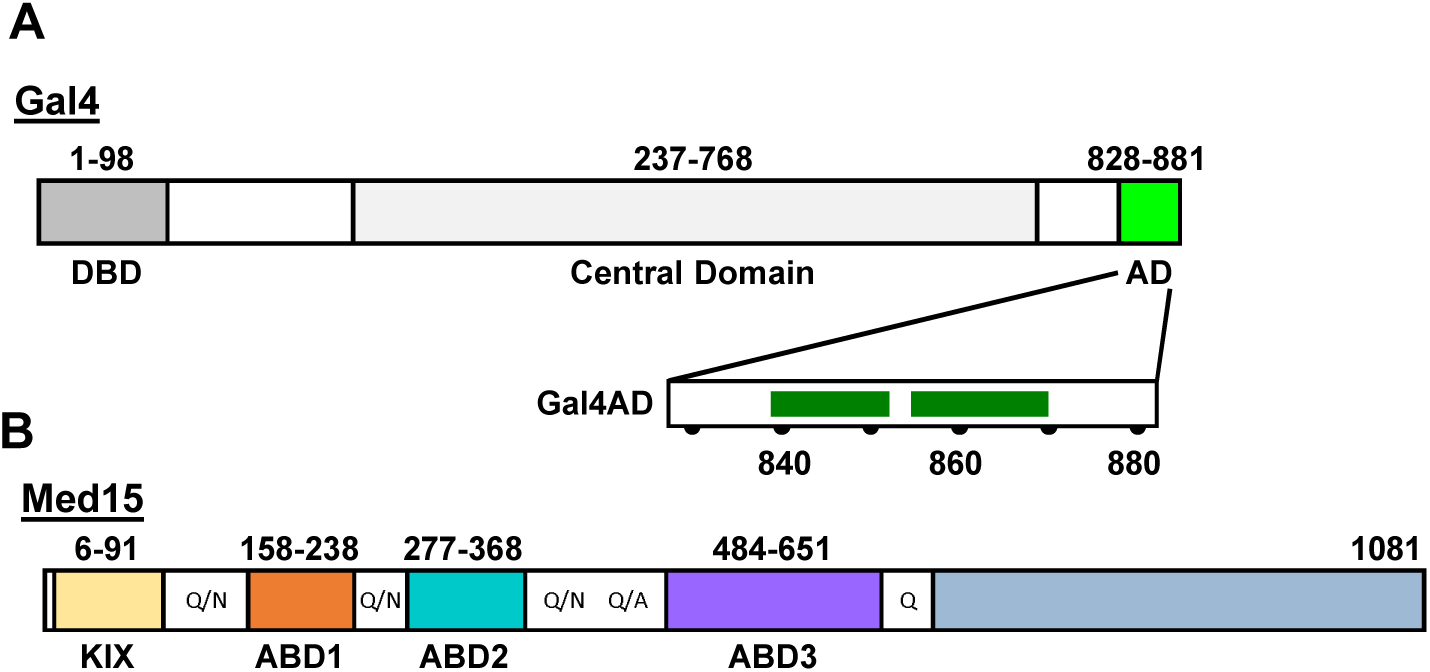
Gal4 AD is an IDR that interacts with Gal15 ABDs. **(A)** Gal4 consists of a DNA binding and dimerization domain (DBD), a Central Domain (CD), and a primary Activator Domain (AD). The expanded view of the AD highlights key hydrophobic Regions 1 and 2 (residues 840-851 and 855-869, respectively). The DBD is structured upon DNA binding; the CD is sequence predicted to be predominantly helical; the AD is intrinsically disordered and includes a Gal80 binding site. **(B)** Med15 consists of four helical activator-binding domains (ABDs) and a C-terminal domain (blue) that is required for incorporation into the Mediator complex. The structured Med15 domains are connected by Q-rich linkers as shown.

Deletion of either the first or last 10 residues of the 828-881 polypeptide had little or no effect on AD function, identifying the minimal Gal4 AD as residues 840 – 871 (Fig. 2). The minimal AD is composed of two fairly short regions, termed Region 1 and Region 2, that each contain large hydrophobic/aromatic residues. Truncations from residue 840 onwards that eliminate part or all of Region 1 reduce transcription activity by ~2-5-fold (847-881 and 852-881, respectively). Truncation from the C-terminal end of the Gal4 AD to eliminate most of Region 2 (840-860) reduced transcription activity by ~7-fold. Thus, the two individual hydrophobic regions of Gal4 have little function on their own but synergize when combined. Our analysis did not reveal the proposed self-inhibitory region located within the AD (residues 872-881) (Piskacek et al., 2017) as removal of these residues (i.e., the construct 828-871) was not associated with an increase in measured activity.

**Figure 2.**
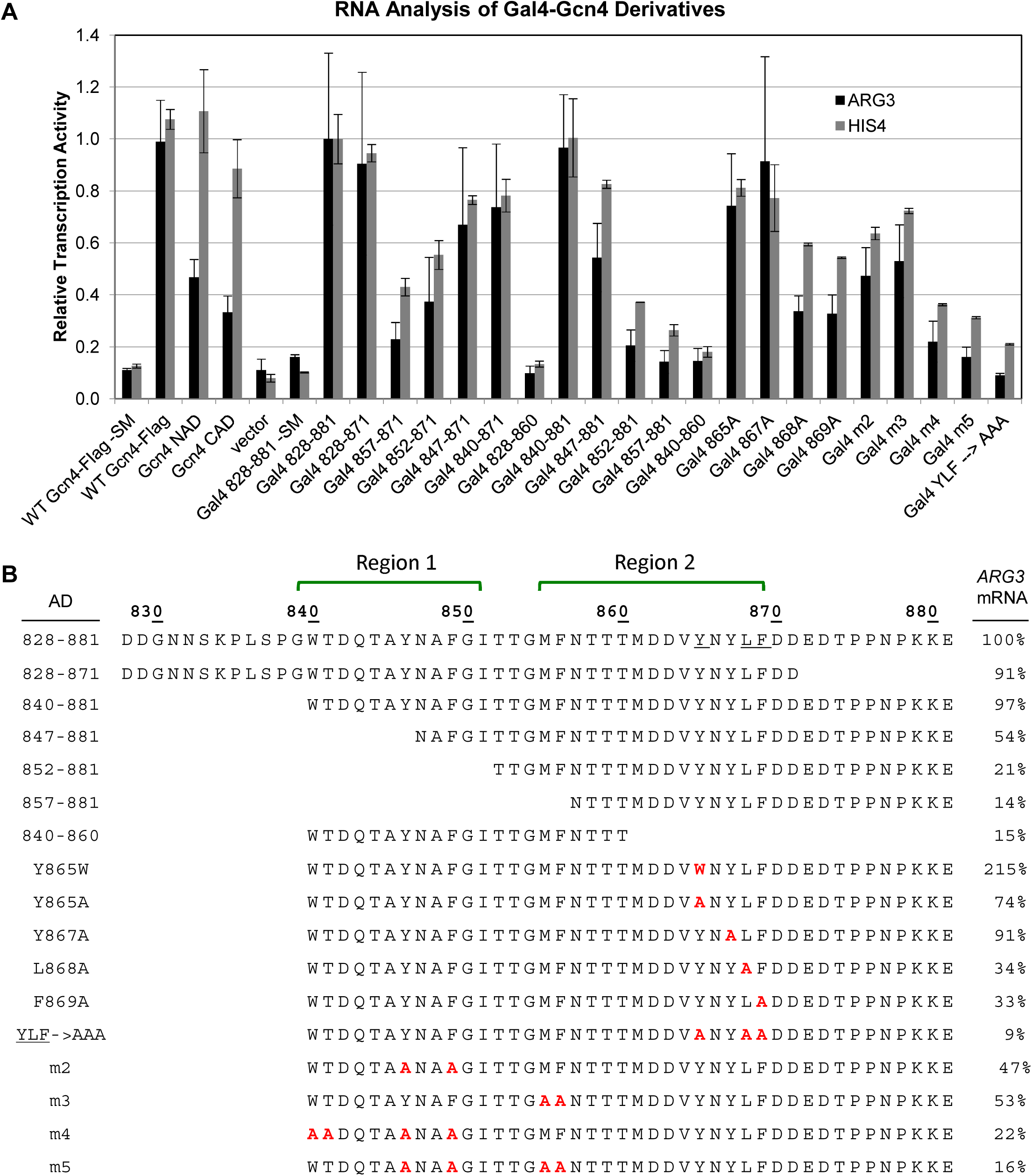
Residues throughout Gal4 840-871 positively contribute to transcription activation. **(A)** RNA analysis of Gal4 AD-Gcn4 DBD derivatives. RNA levels measured by RT qPCR are normalized to *ACT1* mRNA and Gal4 828-881 +SM is set to 1. Gal4 mutants m2-m5 are defined in (B). **(B)** Sequence of Gal4-Gcn4 derivatives and summary of activation assays from (A) at *ARG3*. Gal4 AD hydrophobic patches (green brackets above the sequence) and alanine mutations within these regions (red residues) are shown.

Consistent with the importance of hydrophobicity, substitution of the three aromatic residues in Region 1 to alanine is nearly equivalent to a construct lacking Region 1 (“m4” in Fig. 2). Region 2 contains four aromatic residues, Phe856, Tyr865, Tyr867, and Phe869. With the exception of Phe856, the aromatics are contained within a sequence (^865^YNYLF) that resembles a short sequence motif in the Gcn4 central AD that is critical for function (WXXLF) (Warfield et al., 2014). Despite this similarity, we previously found that shortening Gal4 AD to contain only this segment (857-881) resulted in a construct with little inherent function (See Fig. S1B,C in (Warfield et al., 2014)). Individual alanine substitution of residues in Region 2 have differential effects: replacement of Tyr867 or Tyr865 have only modest effects, while replacement of Phe869 or Leu868 reduced activity by 3-fold. Notably, simultaneous replacement of the three positions that define the motif (i.e., Tyr865, Leu868, and Phe869; “YLF-to-AAA”) produces a construct with at least 10-fold decreased activation of *ARG3*. Altogether our analysis identifies a minimal Gal4 AD (residues 840 – 871) composed of two hydrophobic regions that contribute equally to transcriptional activation of *ARG3*. The more C-terminal region, Region 2, contains a sequence motif similar to that identified in the AD of Gcn4 that is critical to its function. Notably, Region 2 is essentially the same sequence shown to bind to the repressor protein Gal80 with high affinity (Ansari et al., 1998).

### Gal4 AD interacts with Med15 ABDs

Transcriptional activation by Gal4 requires its direct interaction with the Med15 subunit of the Mediator complex. Med15 contains three activator-binding domains, known as ABD1, ABD2, and ABD3 (Fig. 1B) (Herbig et al., 2010; Jedidi et al., 2010). We measured binding affinities of Gal4 AD (construct containing Gal4 residues 828-881 was used for all binding and NMR experiments) with each individual Med15 ABD, KIX, and with a combined ABD123 that contains the three Med15 ABDs connected by shortened linkers. Fluorescence polarization (FP) and/or isothermal calorimetry (ITC) measurements reveal that Gal4 AD binds each ABD with micromolar affinity (Table 1), with strongest binding to ABD1 (K_d_ of 9 μM), followed by ABD3 (K_d_ of 16 μM) and ABD2 (K_d_ of 35 μM). Binding is enhanced in the presence of all three ABDs, with a binding constant value of 2 μM for ABD123. Binding of Gal4 AD to KIX was too weak to measure by FP and ITC. These observations are consistent with other Med15-dependent transcription factors (e.g. Gcn4, Ino2, and Met4) (Pacheco et al., 2018; Tuttle et al., 2018), whose ADs also display micromolar affinities for Med15 and a stronger than additive gain in affinity when multiple ABDs are present (e.g. Gcn4 ADs in Table 1 and (Pacheco et al., 2018)). As well, the rank order of binding strengths for Med15 ABDs to the Gal4 AD and Gcn4 AD are the same, with strongest binding to ABD1 and weakest binding to ABD2, suggesting that the differences in affinity are intrinsic to each ABD. Furthermore, we note that the highest affinity Gal4-Med15 interaction is still ~120-fold lower affinity than the binding of Gal4 to Gal80 repressor (Wu et al., 1996).

**Table 1.** Affinity of ADs for Med15 activator-binding domains

| **Table 1. Affinity of ADs for Med15 activator-binding domains** |  |  |  |
| --- | --- | --- | --- |
| **Binding Data by FP or (ITC)** |  |  |  |
| **Med15** | **AD** | **K<sub>d</sub> (μM)** | **Reference** |
| ABD1 | Gal4 AD | 9.24 ± 1.35, (9.2 ± 0.2) |  |
|  | Gcn4 nAD | 3.3 ± 0.5 | (Herbig et al., 2010) |
|  | Gcn4 cAD | 2.5 ± 0.4 | (Herbig et al., 2010) |
|  | Gcn4 tAD | 2.66 ± 0.22 | (Tuttle et al., 2018) |
| ABD2 | Gal4 AD<sup>1</sup> | 98 ± 15, (35 ± 6.1) |  |
|  | Gcn4 nAD | 21.8 ± 3.8 | (Herbig et al., 2010) |
|  | Gcn4 cAD | 147 ± 30 | (Herbig et al., 2010) |
|  | Gcn4 tAD | 19.5 ± 2.74 | (Tuttle et al., 2018) |
| ABD3 | Gal4 AD | 16.2 ± 1.0 |  |
|  | Gcn4 nAD | 2.6 ± 0.4 | (Herbig et al., 2010) |
|  | Gcn4 cAD | 13.9 ± 0.5 | (Tuttle et al., 2018) |
|  | Gcn4 tAD | 4.40 ± 0.49 | (Tuttle et al., 2018) |
| KIX | Gal4 AD | Unable to measure |  |
|  | Gcn4 nAD | Unable to measure | (Herbig et al., 2010) |
|  | Gcn4 cAD | Unable to measure | (Herbig et al., 2010) |
|  | Gcn4 tAD | Unable to measure | (Tuttle et al., 2018) |
| ABD123 | Gal4 AD | 2.04 ± 0.22 |  |
|  | Gcn4 nAD | 0.36 ± 0.02 | (Herbig et al., 2010) |
|  | Gcn4 cAD | 1.7 ± 0.1 | (Herbig et al., 2010) |
|  | Gcn4 tAD | 0.109 ± 0.012 | (Tuttle et al., 2018) |
| <sup>1</sup>The Gal4 AD:ABD2 K<sub>d</sub> determined by ITC is used for saturation calculations and is consistent with estimates from NMR titrations. |  |  |  |

### Identification of Gal4 AD residues that interact with Med15 ABD1

We used NMR to investigate the mechanism of Gal4-Med15 binding and the role of the Gal4 hydrophobic patches. The Gal4 828-881 polypeptide is intrinsically disordered as evidenced by its narrow ^1^H dispersion in the (^1^H,^15^N)-HSQC NMR spectrum (black peaks, Fig. 3A) and low, uniform T_1_/T_2_ values (green line, Fig. 3C). Despite the narrow spectral dispersion, the NMR spectrum of Gal4 AD could be assigned using conventional triple-resonance protocols. The assignments provided further confirmation that Gal4 AD is disordered, as the backbone ^13^C chemical shifts are close to those predicted for random coil (green bars, Fig. 3D).

**Figure 3.**
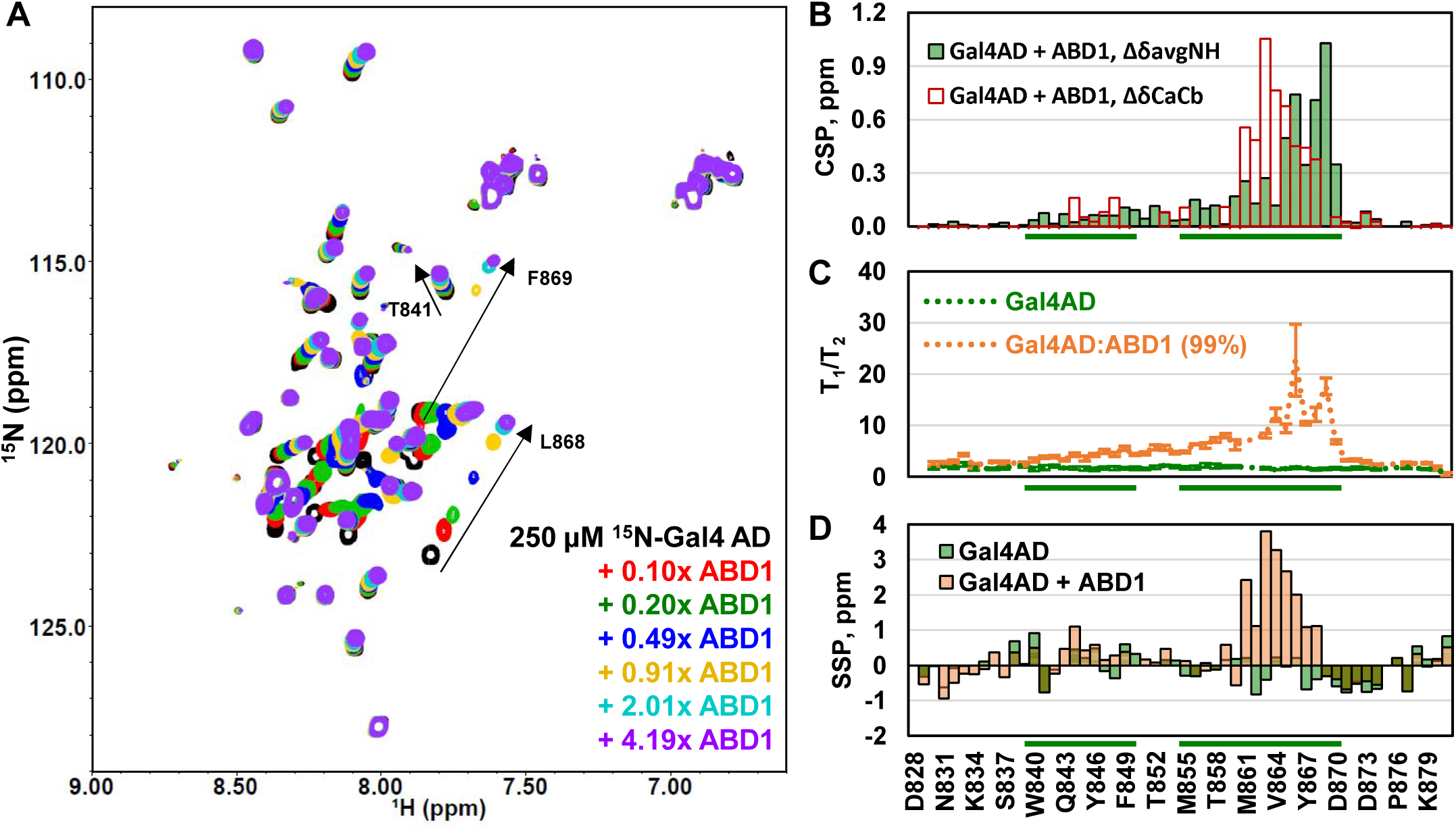
Gal4 AD has a primary region of ABD1 interaction with smaller effects throughout the N-terminus. **(A)** (^1^H,^15^N)-HSQC titration spectra of ^15^N-Gal4 AD with ABD1. Fraction saturations based on the K_d_ are 0 (black), 0.1 (red), 0.19 (green), 0.46 (blue), 0.78 (yellow), 0.98 (cyan), and 0.99 (purple). **(B)** Average amide Chemical Shift Perturbation (CSP) quantifying the free and 99% bound shifts in (A) are shown as green bars. The scaled CSPs for Ca-Cb for free versus bound Gal4 AD are shown as red bars. **(C)** Backbone dynamics of Gal4 AD free (green) and bound to ABD1 (orange). The increase of T_1_/T_2_ in Gal4 AD:ABD1 extending from the primary interacting region to the N-term suggests a weaker interacting role for those regions. **(D)** Secondary Structure Prediction (SSP) for free and bound Gal4 AD. The primary interacting region becomes helical (values towards 4 ppm are consistent with fully helical structure, larger negative values would indicate beta strands). Green bars along the x-axis correspond to Gal4 AD hydrophobic regions of Fig. 1A.

We chose to characterize the effects of Med15 ABD1, the strongest binding ABD, on Gal4 AD. A change in NMR peak positions (chemical shift perturbations; CSPs) indicates a change in chemical environment of the group that gives rise to the NMR peak. As shown in Figure 3A, addition of non-isotopically-labeled Med15 ABD1 results in large changes in peak position (CSP) for some peaks and no perturbation to others. Affected peaks show continuous, fast-exchange chemical shift perturbations as a function of increasing Med15 ABD1. The fast exchange behavior and large CSPs provide an estimate of the exchange rate to be at least 250 s^−1^ and, therefore, a lifetime for the bound state of no more than 4 ms consistent with the micromolar affinity measured for this interaction. Plotting CSPs along the Gal4 AD sequence reveals substantial perturbations to Gal4 residues spanning from 840-870, while the N- and C-termini of the Gal4 AD exhibit only very small CSPs (Fig. 3B) and unchanged dynamics (T_1_/T_2_) (Fig. 3C). Thus, the minimal AD as identified from activity assays (Fig. 2) is congruent with the minimal region affected by ABD1 according to NMR measurements. The largest perturbations are within Region 2, specifically, residues 861-869. Analysis of the backbone chemical shifts for the Gal4 AD/ABD1 complex indicate that residues 861-869 adopt helical structure upon binding to ABD1 (Fig. 3D). This region also shows the largest increase in T_1_/T_2_ dynamics, which is an approximate measure of molecular tumbling time and thus of protein complex size; regions of AD that spend more time bound to ABD1 are expected to have elevated T_1_/T_2_ values. Residues within Region 1 also experience CSPs and small but significant increases in T_1_/T_2_ values, consistent with the region’s contribution to Gal4 AD function. Secondary structure prediction based on backbone chemical shifts indicate that Region 1 residues Gln843 – Phe850 adopt a modest amount of helical structure. Altogether, the data are consistent with transient interactions between ABD1 and Gal4 that are dominated by Gal4 residues 861-869 with lesser contribution from Region 1. Notably, the sequence that appears to dominate the interaction with Med15 ABD1 is the most highly conserved among Gal4 species (**Fig. S1**) and overlaps with the Gal80 binding site.

### Sequence-independent Med15 ABD1 and ABD2 interactions with acidic ADs

To observe the interaction from the perspective of Med15, unlabeled Gal4 AD was titrated into ^15^N-ABD1 or ^15^N-ABD2. As in the reciprocal titrations described above, NMR peaks shifted in fast exchange with increasing Gal4 AD. Remarkably, the chemical shift trajectories of peaks for both ABD1 and ABD2 upon addition of Gal4 AD were nearly identical to those observed upon titration with the tandem Gcn4 ADs (Fig. 4A and **Fig. S2**). Gal4 and Gcn4 have very different primary sequence, but both have numerous hydrophobic regions set in an acidic, disordered background. It is these characteristics that promote binding to the Med15 ABDs and the highly similar perturbations observed point to sequence-independent interactions between the acidic activators and Med15.

**Figure 4.**
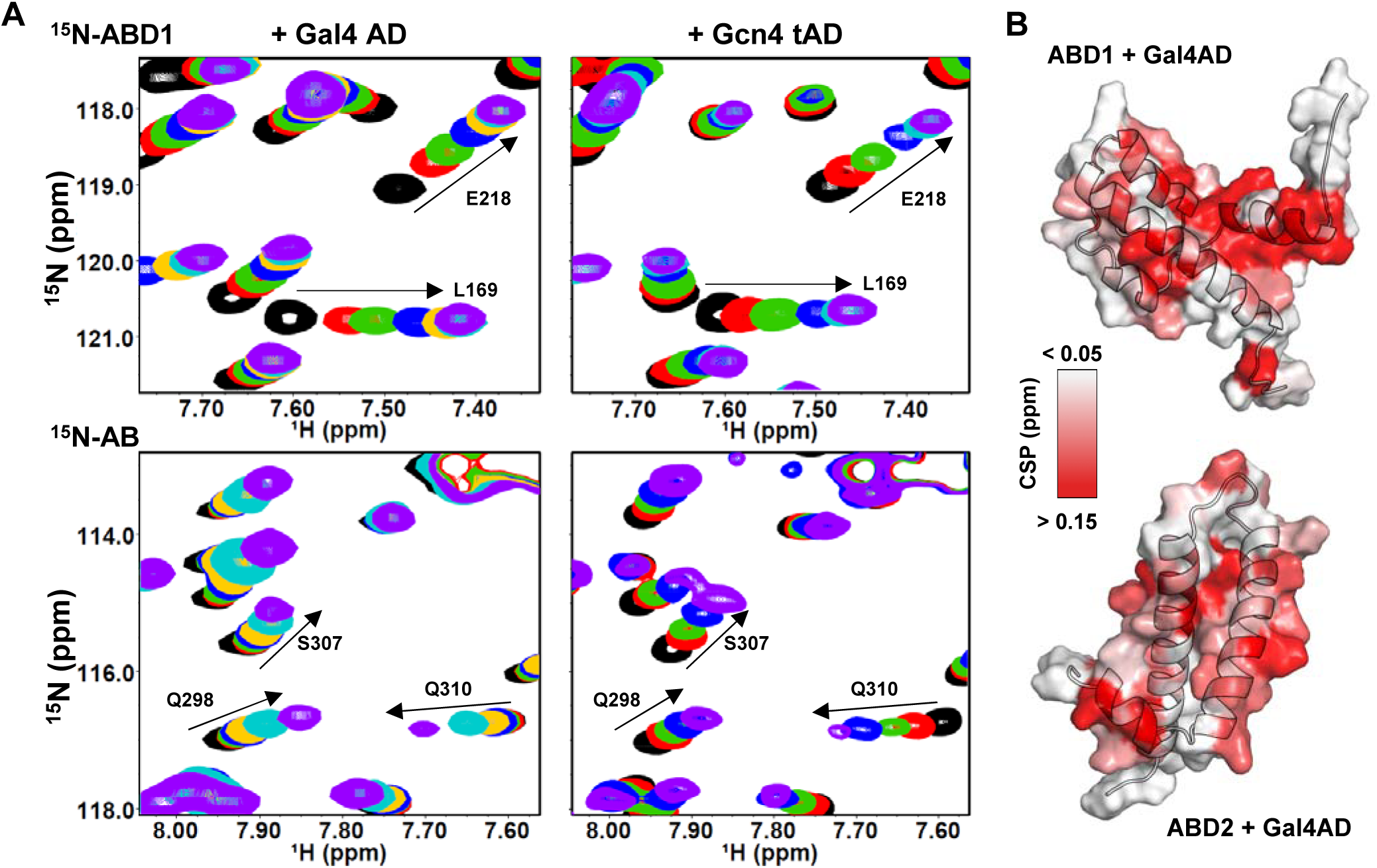
Med15 ABDs have widespread CSPs upon addition of Gal4 AD that resemble the Gcn4 tAD interactions. **(A)** (^1^H,^15^N)-HSQC titration spectra of ^15^N-ABD1 (top) or ^15^N-ABD2 (bottom) plus Gal4 AD (left) or Gcn4 tAD (right). ABD1 and ABD2 show similar chemical shift trajectories regardless of the added activator domain. **(B)** CSPs of ABD + Gal4 AD are plotted on the structures of ABD1 (pdb 2LPB) and ABD2 (pdb 6ALY) with darker red indicating a larger CSP. Disordered ends of ABD1 and ABD2 structures are not shown for clarity. See also **Figure S3**.

Both ABD1 and ABD2 show widespread CSPs upon addition of AD, as plotted on the respective structures in Figure 4B and **Figure S3A**. One face of ABD1 contains a shallow hydrophobic groove that serves as the AD binding site, while binding to ABD2 is distributed over its surface centered on two hydrophobic surface patches. This widespread pattern of CSP is highly similar to that observed with the Gcn4 ADs and is consistent with the formation of fuzzy complexes between Gal4 AD and Med15 (Brzovic et al., 2011; Tuttle et al., 2018; Warfield et al., 2014).

### Gal4 AD forms a fuzzy complex with Med15 ABD1 and ABD2

Fuzzy interactions lack an orientational preference between binding partners. To detect the orientation of Gal4 AD when bound to Med15, we inserted paramagnetic spin-labels into the Gal4 AD sequence. Single Cys mutations for spin-label attachment were placed on either side of the most perturbed and highly conserved Gal4 AD region (Fig. 5A). TEMPO spin-label was attached to either Gal4 AD T860C or T874C and added to ^13^C,^15^N-ABD1 or ^13^C,^15^N-ABD2. The paramagnetic relaxation enhancement (PRE) was determined by quantifying (^1^H,^15^N)- and (^1^H,^13^C)-HSQC spectra with active or inactive spin-label to determine regions of the ABD in proximity to the spin-label site, as peak intensity is lost as a function of time-averaged distance to the spin-label. Previously published results from Gcn4 tAD T82C are included for comparison (Fig. 5B-E) (Tuttle et al., 2018).

**Figure 5.**
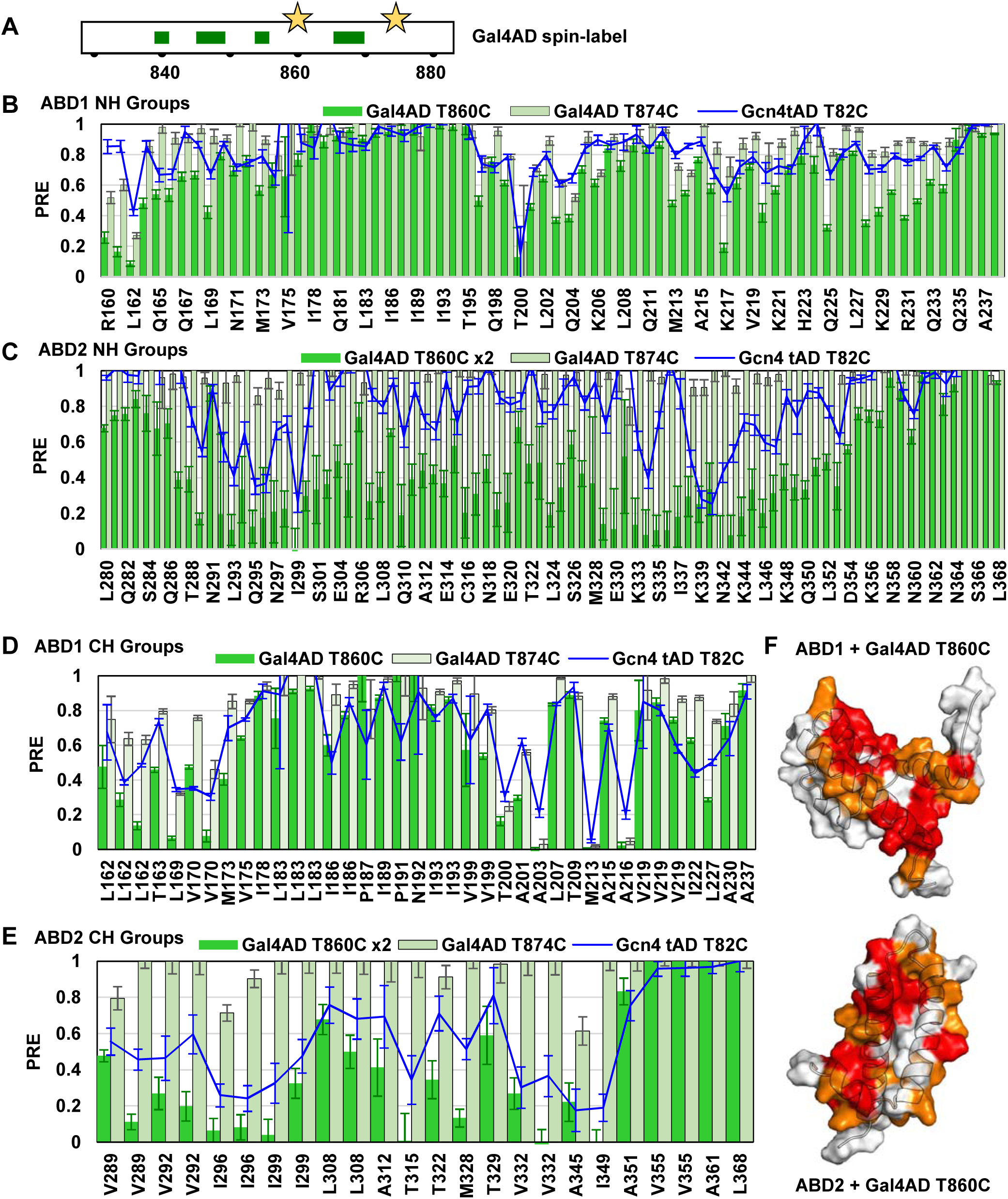
Gcn4 tAD and Gal4 AD spin-labels similarly affect ABD1 and ABD2. (A) Tempo spin-label was attached at either Gal4 AD position T60C or T874C. Paramagnetic Relaxation Effect (PRE) of each spin-labeled AD are shown for NH and CH groups for (**B, D**) ABD1 and (**C, E**) ABD2. Gcn4 tAD T82C spin-label results are included for comparison (Tuttle et al., 2018). The magnitude of the PRE depends on saturation levels of each AD:ABD complex; Gal4 AD T860C has been scaled 2x for ABD2 results since this complex was at higher saturation than the comparison experiments. (**F**) Results of the ABD + Gal4 AD T860C spin-label experiments are plotted on the structures of ABD1 (pdb 2LPB) and ABD2 (pdb 6ALY). The top 25% effected residues are red, the next 25% are orange. See also **Figure S3**.

For ABD1, both Gal4 AD spin-label sites give rise to PREs. However, Gal4 AD T874C gave somewhat smaller PREs overall compared with T860C, despite similar saturation (~60-70%) for both Gal4 AD spin-label samples (Fig. 5B,D). For each spin-labeled AD, loss of ABD1 peak intensity was widespread around the shallow hydrophobic groove where AD binding occurs, with the backside surface largely unaffected (Fig. 5F, **Fig. S3B**). For ABD2, only spin-labeled Gal4 AD T860C gave rise to widespread, significant PREs (Fig. 5C,E). While the ABD2/Gal4 AD T860C spin-labeled sample was more saturated than the T874C spin-label sample (~90% versus ~50% saturation, respectively), this is not expected to be a major reason for the discrepancy, as we have routinely collected our PRE data at around 50% saturation. Instead, it highlights the difference in binding interfaces of ABD1 versus ABD2. Gal4 AD T860 is adjacent to the primarily affected residues 861-870 and has small but significant CSPs upon addition of ABD1 (Fig. 3). In contrast, T874 is located within the Gal4 AD C-terminal residues that do not contribute to transcription activity (Fig. 2) and do not show significant perturbations in the NMR experiments (Fig. 3). The ABD1 binding interface is a shallow hydrophobic groove, which may place Gal4 AD T874 closer on average to ABD1 than for ABD2, which does not have a clear binding pocket. Our results are consistent with a Gal4 AD C-terminal tail that is oriented away from the ABDs and does not contribute to binding. The most affected ABD2 residues in the T874C spin-label experiment are those that are most effected in the T860C experiment (Q295, I296, N297, V332, and A345) and these residues are far apart in the ABD2 structure. The widespread PREs for ABD1 and ABD2 with both Gal4 AD spin-labels are consistent with the results from Gcn4 tAD spin-label experiments, and the pattern of Gal4 AD PREs are very similar to the pattern of Gcn4 tAD PREs (blue lines, Fig. 5). In summary, like with Gcn4-Med15 binding, a single orientation of the bound Gal4 AD cannot satisfy the PRE data and this strongly suggests that Gal4 AD adopts a fuzzy interface upon binding to Med15 ABD1 and ABD2.

## DISCUSSION

Our functional studies identify two hydrophobic-containing regions in a minimal Gal4 AD that contribute to transcriptional activation. Both regions are required for robust transcription activation in vivo, as each individual region has only modest activation activity on its own. Such behavior is reminiscent of the Gcn4 N-terminal AD (nAD), in which four hydrophobic clusters distributed over 100 residues are required for full function and no single hydrophobic cluster can activate transcription effectively on its own. Both Gal4 AD regions interact directly with ABDs of Med15, as evidenced by NMR, with Region 2 appearing to dominate the binding. We note that the most C-terminal hydrophobic cluster in the Gcn4 nAD also shows the largest chemical shift perturbations, despite additional clusters being functionally important. These two examples contrast with the Gcn4 cAD, whose function is largely encoded within a single short sequence embedded in a disordered, acidic polypeptide (Brzovic et al., 2011; Staller et al., 2018; Warfield et al., 2014). Nevertheless, important hydrophobic residues are also spread over a 20-30 residue range in yeast transcription activators Met4 and Ino2, suggesting that hydrophobic residues distributed over an AD are more common than a single short dominant sequence. This model offers a rationale for why conserved sequence motif(s) have not been identified among transcription activators. Furthermore, the interactions are weak and transient, with an upper limit on the lifetime of an AD/ABD1 complex of around 4 ms. We suggest that the multiple short hydrophobic regions contribute to Med15 binding by a mechanism similar to avidity or allovalency (Olsen et al., 2017). By this mechanism, multiple weak binding sites in the AD act to increase the effective affinity for multiple binding sites on Med15 by decreasing the dissociation rate of the two molecules and effectively increasing the local concentration of the binding partners. This binding mechanism can also explain the positive contribution of entropy to Med15 binding observed with Gcn4, Ino2, Met4, and Gal4 ADs (Brzovic et al., 2011; Pacheco et al., 2018; Warfield et al., 2014).

The NMR results confirm that the Gal4 AD is disordered in its unbound state and that part of Region 2 (residues 861-868) adopts helical structure upon binding (Fig. 3D). Yet despite adopting structure, this hydrophobic region binds to ABD1 without orientational preference. The two Med15 ABDs investigated in this study are each structured on their own and present preformed binding surfaces. The remarkable similarities of the CSPs induced by Gal4 AD and Gcn4 tAD, which have very different sequences, on each of the ABDs indicate that the two ADs interact similarly with the ABD surfaces, via a hydrophobic cloud rather than via defined sidechain-to-sidechain interactions. Such sequence-independent effects, along with the lack of orientational binding preference, are hallmarks of fuzzy binding. Transient, dynamic interactions of multiple regions of Gal4 AD with each of the Med15 ABDs is consistent with a fuzzy free-for-all mechanism (Fig. 6) (Tuttle et al., 2018).

**Figure 6.**
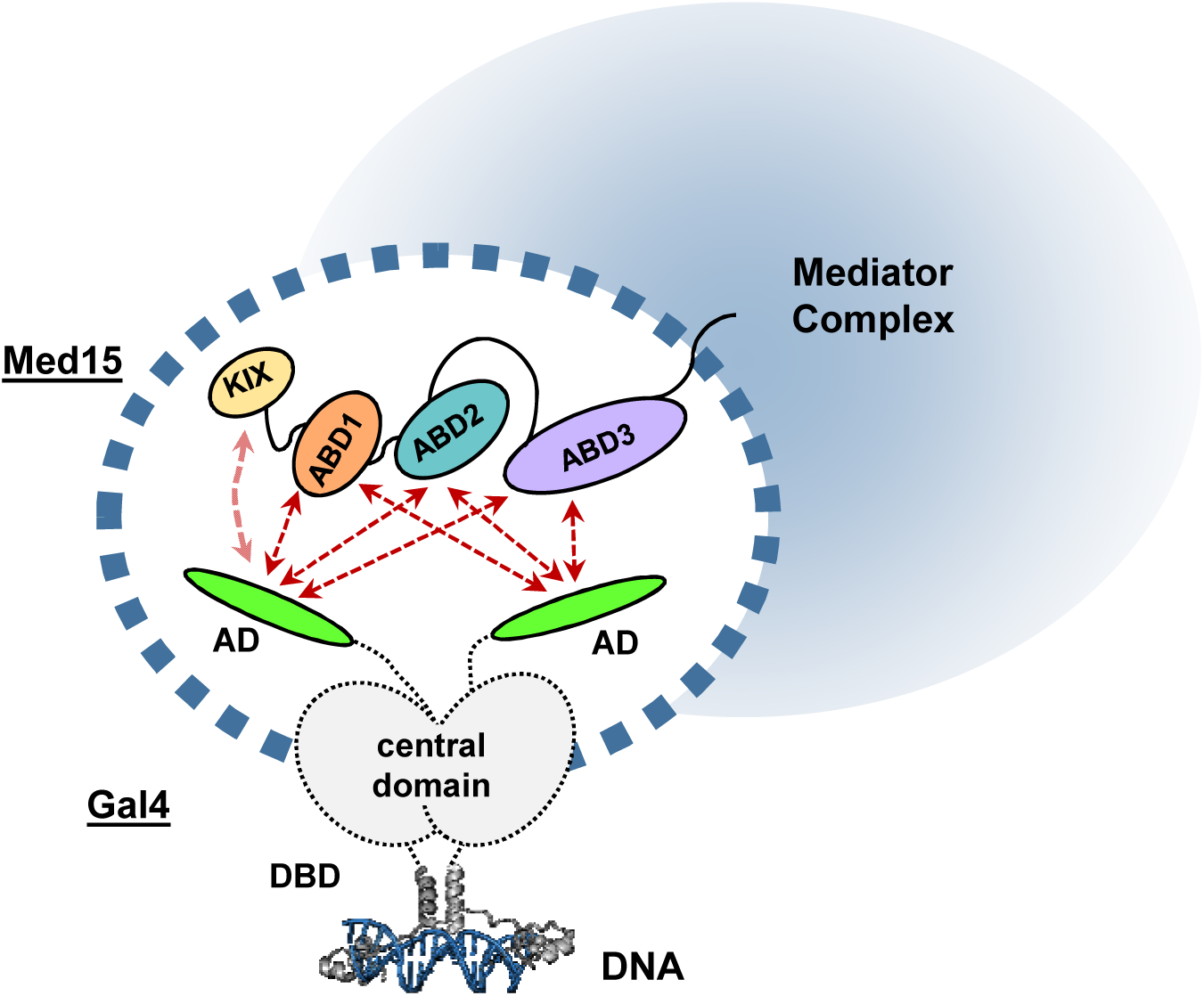
Model for interaction of the Gal4 AD and Med15 ABDs in a dynamic fuzzy complex. Although the Gal4 AD sequence is very different from the Gcn4 ADs and tightly binds Gal80 repressor, the two activators bind Med15 using the same fuzzy free-for-all mechanism. Gal4DBD:DNA structure is from pdb 1D66.

Our results lead to two general conclusions. First, we propose that the fuzzy-free-for-all mechanism is common among transcription factors with acidic activation domains. This conclusion is based on the demonstrated similarities between the ways in which Gal4 and Gcn4 engage Med15, despite their lack of sequence conservation and the very different mode of regulation of Gal4 and Gnc4 transcription factors. The notion of a common mode of action among ADs is non-trivial. Although the ADs of different transcription factors can be interchangeable (Hope and Struhl, 1986; Sadowski et al., 1988), each intrinsically disordered AD has undergone specific positive selection (Afanasyeva et al., 2018). For example, the Gal4 AD has evolved to bind Gal80 repressor with high affinity and sequence specificity yet the same sequence is directly involved in sequence-independent fuzzy interaction with Med15. In contrast, Gcn4 is under no analogous constraint.

A second conclusion that can be drawn from our study is that the mode of binding of intrinsically disordered ADs is dictated predominantly by the binding partner, rather than by the disordered component. This conclusion is supported by two observations. First, the binding preferences/strengths of the ADs from Gal4 and Gcn4 to the three ABDs of Med15 follow the same trend, with highest affinity for ABD1 and lowest affinity for ABD2. What is in common are the ABD structures, not the ADs. Second, within the Gal4 AD, an identical sequence (contained within Region 2) binds as a fuzzy complex to Med15 ABDs but binds as a well-oriented, structured polypeptide to the Gal80 repressor (Kumar et al., 2008; Thoden et al., 2008). This is strong evidence that it is the binding partner that dictates whether a disordered ligand will remain disordered while bound, will adopt structure but bind in a fuzzy manner, or will bind in a canonical, orientationally-defined manner.

Questions remain regarding the overall mechanism of transcription activation. For systems with dynamic weak interactions, condensate formation is proposed to help compartmentalize the components and thereby decrease the frequency of non-productive encounters, acting to increase local concentrations of components and drive the frequency of these interactions (Liu and Tjian, 2018). Recent studies demonstrate the importance of condensate formation for transcription activation via super enhancers, with the AD appearing to be an important driving force for condensate formation (Boija et al., 2018; Hnisz et al., 2017; Sabari et al., 2018; Shrinivas et al., 2019). A fuzzy free-for-all binding mechanism can be consistent with the formation of condensates (Hahn, 2018). Furthermore, the multiple transcription factor binding sites that often occur at promoter regions can contribute to higher local concentrations. For example, the *HIS3* promoter has seven Gcn4 consensus binding regions; *HIS5* promoter contains five potential sites for Gcn4 (Arndt and Fink, 1986); and the promoter controlling *GAL1*, *GAL4*, and *GAL10* transcription contains four Gal4 DBD sites to which Gal4 binds cooperatively (Ma and Ptashne, 1987b). Each DNA site binds a dimer of the relevant transcription factor, presenting two ADs, each of which contain multiple hydrophobic binding regions per site. In turn, the ADs could simultaneously interact with components of Mediator or other coactivator binding partners via a fuzzy free-for-all mechanism to promote transcription activation (Fig. 6). It is as yet unknown whether condensate formation is a direct consequence of such interactions or whether such formations are functionally relevant. Nevertheless, ADs have been implicated in formation of condensates and there may be further dependence on the Q-rich linker regions of the coactivators (Alberti, 2017; Boija et al., 2018; Cooper and Fassler, 2019; Sabari et al., 2018).

Taken together, our conclusions indicate that transcription activation by Gal4 and Gcn4 with Med15 occurs via a common interaction mechanism that is dictated by the properties of the Med15 ABDs. A remaining question is whether these same ADs use a fuzzy binding mechanism in their interactions with other coactivators such as with subunits of SAGA, or whether the Med15 ABDs are uniquely configured to facilitate fuzzy interactions.

## EXPERIMENTAL PROCEDURES

Further details and an outline of resources used in this work can be found in Supplemental Experimental Procedures.

## CONTACT FOR REAGENT AND RESOURCE SHARING

Further information and requests for resources and reagents should be directed to and will be fulfilled by the Lead Contacts, Steve Hahn or Rachel Klevit.

### Yeast strains and plasmids

Yeast strain SHY944 (*matα ade2Δ::hisG his3Δ200 leu2Δ0 lys2Δ0 met15Δ0 trp1Δ63 ura3Δ0 gcn4Δ::KanMx gal80Δ::HIS3*) was transformed with the indicated Gcn4 or Gal4AD-Gcn4 containing plasmids that are derivatives of the *LEU2*-containing vector pRS315. All Gcn4 derivatives contained a C-terminal 3X-Flag tag.

### mRNA analysis

Yeast strains were grown in duplicate at 30°C to an optical density at 600 nm (OD600) of 0.5 to 0.8 in 2% (wt/vol) dextrose synthetic complete media lacking Ile, Val and Leu. Cells were induced with 0.5 μg/ml SM for 90 min to induce amino acid starvation (27), RNA was extracted and assayed in triplicate by reverse transcription-quantitative PCR (RT-qPCR), and the results were analyzed as described previously (Herbig et al., 2010). Data are plotted in Fig2 as mRNA ratios of *ARG3* to *ACT1* or *HIS4* to *ACT1*.

### Protein purification

All proteins were expressed in BL21 (DE3) RIL E. coli. Med15 484-651 (ABD3) and Med15 158-651 Δ239-272, Δ373-483 (ABD123) were expressed as N-terminal His6-tagged proteins. All other Med15 and Gal4 constructs were expressed as N-terminal His6-SUMO-tagged proteins. Cells were lysed in 50 mM HEPES pH 7.0, 500 mM NaCl, 40 mM Imidazole, 10% glycerol, 1 mM PMSF, 5 mM DTT and purified using Ni-Sepharose High Performance resin (GE Healthcare). Proteins were eluted in 50 mM HEPES pH 7.0, 500 mM NaCl, 500 mM Imidazole, 10% glycerol, 1 mM PMSF, 1 mM DTT. Purified SUMO-tagged proteins were concentrated using 10K MW cutoff centrifugal filters (Millipore), diluted 10x in 50 mM HEPES pH 7.0, 500 mM NaCl, 40 mM Imidazole, 10% glycerol, 1 mM PMSF, 5 mM DTT, and digested with SUMO protease for 3-5 hrs at room temperature using ~1:800 protease:protein ratio. Cleaved His6-Sumo tag was removed using Ni-Sepharose. Med15 polypeptides were further purified using HiTrap Heparin (GE Healthcare) in 20 mM HEPES pH 7, 1 mM DTT, 0.5mM PMSF eluting with either a 50-350 mM NaCl gradient (ABD1 and ABD2) or a 200-600 mM NaCl gradient (ABD3 and ABD123). Gal4 AD was further purified by chromatography on Source 15Q (GE Healthcare) using a 50-350 mM NaCl gradient. All proteins were further purified using size exclusion chromatography on Superdex 75 10/30 (GE Healthcare). Proteins used in fluorescence polarization and ITC were eluted in 20 mM KH_2_PO_4_, pH 7.5, 200 mM KCl. Proteins used in NMR were eluted in 20 mM NaH_2_PO_4_ pH 6.5, 200 mM NaCl, 0.1 mM EDTA, 0.1 mM PMSF, 5 mM DTT. The concentration of the purified proteins was determined by UV/Vis spectroscopy with extinction coefficients calculated with ProtParam (Gasteiger et al., 2005).

### FP and ITC binding experiments

Gal4 peptides used in fluorescence polarization were labeled with Oregon Green 488 dye (Invitrogen) as previously described (Herbig et al., 2010). FP measurements were conducted using a Beacon 2000 instrument as previously described (Herbig et al., 2010). Titrations between Gal4 AD and Med15 were performed with 15 concentrations of Med15 spanning 0-250 μM (KIX, ABD1, ABD2, ABD3, ABD123) at 22 °C. All Gal4 AD vs Med15 titrations were done in triplicate except for ABD123, which was done in duplicate. FP data was analyzed using Prism 7 (Graphpad Software, Inc.) to perform non-linear regression analysis using the one-site total binding model Y=B_max_*X/(K_d_+X) + NS*X + Background where Y equals arbitrary polarization units and X equals Med15 concentration.

Isothermal calorimetry (ITC) titrations were performed using a Microcal ITC200 Microcalorimeter in 20 mM KH2PO4, pH 7.5, 200 mM KCl as described in (Brzovic et al., 2011). The following protein concentrations were used, with the syringe molecule listed first and the cell molecule listed second: Gal4 AD (1.11 mM) vs. Med15 ABD1 (0.102 mM); Gal4 AD (1.11 mM) vs. Med15 ABD2 (0.11 mM). The injection settings were 15 × 2.55 ul with 180 sec spacing, 22 °C cell temp, Ref power = 10 ucal/sec, and 1000 rpm stir speed. Calorimetric data were plotted and fit with a single binding site model using Origin 7.0 software (Microcal).

### NMR experiments and resonance assignments

NMR HSQC titration, dynamics, and spin-label experiments were completed on a Bruker 500 MHz Avance III spectrometer. All spectra were collected at 25 °C in NMR buffer (20 mM sodium phosphate, 150 mM NaCl, 0.1 mM EDTA, 0.1 mM PMSF, and 5 mM DTT, pH 6.5) with 7% D_2_O, unless otherwise specified. Spin-label samples were in NMR buffer but with no DTT.

(^1^H,^15^N)- and (^1^H,^13^C)-HSQC titration experiments were completed for Gal4-AD:Med15-ABD combinations by adding unlabeled Gal4 AD or ABD (ABD1 or ABD2) to a ~200 μM [^13^C,^15^N]-AD or ABD sample, maintaining a constant concentration of the labeled species. T_1_ and T_2_ ^15^N-Trosy HSQC experiments for 200 μM [^13^C,^15^N]-Gal4 AD and 200 μM [^13^C,^15^N]-Gal4 AD + 640 μM ABD1 were collected in an interleaved manner with 8 points each (Zhu et al., 2000). T_1_ delays were 10, 40, 80, 120, 160, 320, 640, and 1000 ms; T_2_ cpmg loop delays were 8.48, 16.96, 25.44, 33.92, 42.40, 50.88, 59.36, and 67.84 ms. T_1_ and T_2_ for each residue were fitted to a single exponential with errors reflecting the quality in the fit.

Paramagnetic relaxation enhancement (PRE) experiments were performed as for Gcn4 and ABD1/2 (Tuttle et al., 2018). The spin-label 4-(2-Iodoacetamido)-TEMPO was incorporated at a single Cys mutant for Gal4 AD T860C and T874C by incubating the AD Cys mutants with 10x TEMPO overnight at room temperature, followed by several hours at 30 °C. Excess TEMPO was removed by elution over a Nap-10 column and buffer exchange during concentration. ^15^N- and ^13^C-HSQC spectra of 250 μM [^13^C,^15^N]-ABD1 and 200 μM [^13^C,^15^N]-ABD2 with spin-labeled Gal4 ADs were collected in the presence (reference intensity, I_dia_) and absence of 3 mM ascorbic acid (I_para_). Spin-label Gal4 AD concentrations were 180 μM for T860C with ABD1, 200 μM T874C with ABD1, 150 μM T860C with ABD2, and 650 μM for Gal4 T860C with ABD2.

Backbone chemical shift assignments were determined from standard backbone triple-resonance experiments (HNCA, HNCOCA, HNCOCACB, HNCACB, and HNCO) obtained on a 560 μM [^13^C,^15^N]-Gal4 AD sample and a 200 μM [^13^C,^15^N]-Gal4 AD + 640 μM ABD1 sample using a Bruker 600 MHz Avance spectrometer with cryoprobe.

The NH chemical shift perturbation was calculated according to Δδ_NH_ (ppm) = sqrt [Δδ_H_^2^ + (Δδ_N_/5)^2^]. Component ^1^H and ^15^N chemical shift differences are (free – bound) in ppm. The CaCb CSP was calculated according to Δδ_CaCb_ (ppm) = 0.25*[(Cα-Cβ)_free_-(Cα-Cβ)_bound_]. Secondary structure propensity was determined from Δδ(Cα-Cβ) = (Cα-Cβ)_measured_ – (Cα-Cβ)_random coil_, where the random coil shifts were generated from Gal4 AD sequence using a webserver (https://www1.bio.ku.dk/english/research/bms/research/sbinlab/groups/mak/randomcoil/script/) (Kjaergaard and Poulsen, 2011).

## QUANTIFICATION AND STATISTICAL ANALYSIS

### mRNA quantification

Steady-state mRNA levels were quantitated by RT qPCR as described above and in (Herbig et al., 2010).

### NMR analysis

All NMR spectra were processed using NMRPipe (Delaglio et al., 1995), and analyzed in nmrViewJ (Johnson, 2004). NMR peak intensities for spin-label experiments were quantified using NMRviewJ and error bars reflect noise levels of the spectra. Percent saturation was calculated based on the K_d_ for each A:B complex according to %sat = [AB]/A = (1/A)*(C/2+(C^2^ – 4*A*B)^0.5^), where A=[A_total_], B=[B_total_], and C=A+B+K_d_.

## DATA AVAILABILITY

NMR chemical shifts have been deposited to the BioMagResBank: Gal4 AD 828-881 (accession number 50086).

### ACKNOWLEDGEMENTS

We thank all members of the Klevit and Hahn laboratories for their suggestions and comments during this work. This work was funded by NIH RO1 GM075114 to SH and REK.

## AUTHOR CONTRIBUTIONS

L.M.T., S.H., and R.E.K designed the research. L.M.T, D.P., L.W., and S.H, performed the experiments. L.M.T., D.P., L.W., S.H, and R.E.K analyzed data. L.M.T., D.P., S.H, and R.E.K. wrote the paper.

## SUPPLEMENTAL ITEMS

**Document S1. Supplemental Experimental Procedures, Figures S1-S3.**

## REFERENCES

Afanasyeva, A., Bockwoldt, M., Cooney, C.R., Heiland, I., and Gossmann, T.I. (2018). Human long intrinsically disordered protein regions are frequent targets of positive selection. Genome research 28, 975–982.

Alberti, S. (2017). The wisdom of crowds: regulating cell function through condensed states of living matter. J Cell Sci 130, 2789–2796.

Ansari, A.Z., Reece, R.J., and Ptashne, M. (1998). A transcriptional activating region with two contrasting modes of protein interaction. Proc Natl Acad Sci U S A 95, 13543–13548.

Arndt, K., and Fink, G.R. (1986). GCN4 protein, a positive transcription factor in yeast, binds general control promoters at all 5’ TGACTC 3’ sequences. Proc Natl Acad Sci U S A 83, 8516–8520.

Boija, A., Klein, I.A., Sabari, B.R., Dall’Agnese, A., Coffey, E.L., Zamudio, A.V., Li, C.H., Shrinivas, K., Manteiga, J.C., Hannett, N.M., et al. (2018). Transcription Factors Activate Genes through the Phase-Separation Capacity of Their Activation Domains. Cell 175, 1842–1855 e1816.

Brzovic, P.S., Heikaus, C.C., Kisselev, L., Vernon, R., Herbig, E., Pacheco, D., Warfield, L., Littlefield, P., Baker, D., Klevit, R.E., et al. (2011). The acidic transcription activator Gcn4 binds the mediator subunit Gal11/Med15 using a simple protein interface forming a fuzzy complex. Mol Cell 44, 942–953.

Cooper, D.G., and Fassler, J.S. (2019). Med15: Glutamine-Rich Mediator Subunit with Potential for Plasticity. Trends Biochem Sci 44, 737–751.

Delaglio, F., Grzesiek, S., Vuister, G.W., Zhu, G., Pfeifer, J., and Bax, A. (1995). NMRPipe: a multidimensional spectral processing system based on UNIX pipes. J Biomol NMR 6, 277–293.

Erkina, T.Y., and Erkine, A.M. (2016). Nucleosome distortion as a possible mechanism of transcription activation domain function. Epigenetics Chromatin 9, 40.

Fischer, J.A., Giniger, E., Maniatis, T., and Ptashne, M. (1988). GAL4 activates transcription in Drosophila. Nature 332, 853–856.

Fishburn, J., Mohibullah, N., and Hahn, S. (2005). Function of a eukaryotic transcription activator during the transcription cycle. Mol Cell 18, 369–378.

Gasteiger, E., Hoogland, C., Gattiker, A., Duvaud, S., Wilkins, M.R., Appel, R.D., and Bairoch, A. (2005). Protein Identification and Analysis Tools on the ExPASy Server. In The Proteomics Protocols Handbook, J.M. Walker, ed. (Humana Press), pp. 571–607

Hahn, S. (2018). Phase Separation, Protein Disorder, and Enhancer Function. Cell 175, 1723–1725.

Hahn, S., and Young, E.T. (2011). Transcriptional regulation in Saccharomyces cerevisiae: transcription factor regulation and function, mechanisms of initiation, and roles of activators and coactivators. Genetics 189, 705–736.

Herbig, E., Warfield, L., Fish, L., Fishburn, J., Knutson, B.A., Moorefield, B., Pacheco, D., and Hahn, S. (2010). Mechanism of Mediator recruitment by tandem Gcn4 activation domains and three Gal11 activator-binding domains. Mol Cell Biol 30, 2376–2390.

Hnisz, D., Shrinivas, K., Young, R.A., Chakraborty, A.K., and Sharp, P.A. (2017). A Phase Separation Model for Transcriptional Control. Cell 169, 13–23.

Hope, I.A., and Struhl, K. (1986). Functional dissection of a eukaryotic transcriptional activator protein, GCN4 of yeast. Cell 46, 885–894.

Jackson, B.M., Drysdale, C.M., Natarajan, K., and Hinnebusch, A.G. (1996). Identification of seven hydrophobic clusters in GCN4 making redundant contributions to transcriptional activation. Mol Cell Biol 16, 5557–5571.

Jedidi, I., Zhang, F., Qiu, H., Stahl, S.J., Palmer, I., Kaufman, J.D., Nadaud, P.S., Mukherjee, S., Wingfield, P.T., Jaroniec, C.P., et al. (2010). Activator Gcn4 employs multiple segments of Med15/Gal11, including the KIX domain, to recruit mediator to target genes in vivo. J Biol Chem 285, 2438–2455.

Jeong, C.J., Yang, S.H., Xie, Y., Zhang, L., Johnston, S.A., and Kodadek, T. (2001). Evidence that Gal11 protein is a target of the Gal4 activation domain in the mediator. Biochemistry 40, 9421–9427.

Johnson, B.A. (2004). Using NMRView to visualize and analyze the NMR spectra of macromolecules. Methods Mol Biol 278, 313–352.

Johnston, S.A., Salmeron, J.M., Jr., and Dincher, S.S. (1987). Interaction of positive and negative regulatory proteins in the galactose regulon of yeast. Cell 50, 143–146.

Kakidani, H., and Ptashne, M. (1988). GAL4 activates gene expression in mammalian cells. Cell 52, 161–167.

Keegan, L., Gill, G., and Ptashne, M. (1986). Separation of DNA binding from the transcription-activating function of a eukaryotic regulatory protein. Science 231, 699–704.

Kjaergaard, M., and Poulsen, F.M. (2011). Sequence correction of random coil chemical shifts: correlation between neighbor correction factors and changes in the Ramachandran distribution. J Biomol NMR 50, 157–165.

Kumar, P.R., Yu, Y., Sternglanz, R., Johnston, S.A., and Joshua-Tor, L. (2008). NADP regulates the yeast GAL induction system. Science 319, 1090–1092.

Lavy, T., Kumar, P.R., He, H., and Joshua-Tor, L. (2012). The Gal3p transducer of the GAL regulon interacts with the Gal80p repressor in its ligand-induced closed conformation. Genes Dev 26, 294–303.

Li, Y., Chen, G., and Liu, W. (2010). Multiple metabolic signals influence GAL gene activation by modulating the interaction of Gal80p with the transcriptional activator Gal4p. Mol Microbiol 78, 414–428.

Liu, Z., and Tjian, R. (2018). Visualizing transcription factor dynamics in living cells. J Cell Biol 217, 1181–1191.

Ma, J., Przibilla, E., Hu, J., Bogorad, L., and Ptashne, M. (1988). Yeast activators stimulate plant gene expression. Nature 334, 631–633.

Ma, J., and Ptashne, M. (1987a). The carboxy-terminal 30 amino acids of GAL4 are recognized by GAL80. Cell 50, 137–142.

Ma, J., and Ptashne, M. (1987b). Deletion analysis of GAL4 defines two transcriptional activating segments. Cell 48, 847–853.

Olsen, J.G., Teilum, K., and Kragelund, B.B. (2017). Behaviour of intrinsically disordered proteins in protein-protein complexes with an emphasis on fuzziness. Cell Mol Life Sci 74, 3175–3183.

Pacheco, D., Warfield, L., Brajcich, M., Robbins, H., Luo, J., Ranish, J., and Hahn, S. (2018). Transcription Activation Domains of the Yeast Factors Met4 and Ino2: Tandem Activation Domains with Properties Similar to the Yeast Gcn4 Activator. Mol Cell Biol 38.

Piskacek, M., Havelka, M., Rezacova, M., and Knight, A. (2017). The 9aaTAD Is Exclusive Activation Domain in Gal4. PLoS One 12, e0169261.

Ptashne, M., and Gann, A. (2002). Genes & signals (Cold Spring Harbor, New York: Cold Spring Harbor Laboratory Press).

Reeves, W.M., and Hahn, S. (2005). Targets of the Gal4 transcription activator in functional transcription complexes. Mol Cell Biol 25, 9092–9102.

Sabari, B.R., Dall’Agnese, A., Boija, A., Klein, I.A., Coffey, E.L., Shrinivas, K., Abraham, B.J., Hannett, N.M., Zamudio, A.V., Manteiga, J.C., et al. (2018). Coactivator condensation at super-enhancers links phase separation and gene control. Science 361.

Sadowski, I., Ma, J., Triezenberg, S., and Ptashne, M. (1988). GAL4-VP16 is an unusually potent transcriptional activator. Nature 335, 563–564.

Salmeron, J.M., Jr., Leuther, K.K., and Johnston, S.A. (1990). GAL4 mutations that separate the transcriptional activation and GAL80-interactive functions of the yeast GAL4 protein. Genetics 125, 21–27.

Shrinivas, K., Sabari, B.R., Coffey, E.L., Klein, I.A., Boija, A., Zamudio, A.V., Schuijers, J., Hannett, N.M., Sharp, P.A., Young, R.A., et al. (2019). Enhancer Features that Drive Formation of Transcriptional Condensates. Mol Cell 75, 549–561 e547.

Sil, A.K., Alam, S., Xin, P., Ma, L., Morgan, M., Lebo, C.M., Woods, M.P., and Hopper, J.E. (1999). The Gal3p-Gal80p-Gal4p transcription switch of yeast: Gal3p destabilizes the Gal80p-Gal4p complex in response to galactose and ATP. Mol Cell Biol 19, 7828–7840.

Sood, V., and Brickner, J.H. (2017). Genetic and Epigenetic Strategies Potentiate Gal4 Activation to Enhance Fitness in Recently Diverged Yeast Species. Curr Biol 27, 3591–3602 e3593.

Staller, M.V., Holehouse, A.S., Swain-Lenz, D., Das, R.K., Pappu, R.V., and Cohen, B.A. (2018). A High-Throughput Mutational Scan of an Intrinsically Disordered Acidic Transcriptional Activation Domain. Cell Syst 6, 444–455 e446.

Stone, G., and Sadowski, I. (1993). GAL4 is regulated by a glucose-responsive functional domain. EMBO J 12, 1375–1385.

Suzuki, Y., Nogi, Y., Abe, A., and Fukasawa, T. (1988). GAL11 protein, an auxiliary transcription activator for genes encoding galactose-metabolizing enzymes in Saccharomyces cerevisiae. Mol Cell Biol 8, 4991–4999.

Thoden, J.B., Ryan, L.A., Reece, R.J., and Holden, H.M. (2008). The interaction between an acidic transcriptional activator and its inhibitor. The molecular basis of Gal4p recognition by Gal80p. J Biol Chem 283, 30266–30272.

Tuttle, L.M., Pacheco, D., Warfield, L., Luo, J., Ranish, J., Hahn, S., and Klevit, R.E. (2018). Gcn4-Mediator Specificity Is Mediated by a Large and Dynamic Fuzzy Protein-Protein Complex. Cell Rep 22, 3251–3264.

Warfield, L., Tuttle, L.M., Pacheco, D., Klevit, R.E., and Hahn, S. (2014). A sequence-specific transcription activator motif and powerful synthetic variants that bind Mediator using a fuzzy protein interface. Proc Natl Acad Sci U S A 111, E3506–3513.

Webster, N., Jin, J.R., Green, S., Hollis, M., and Chambon, P. (1988). The Yeast Uasg Is a Transcriptional Enhancer in Human Hela-Cells in the Presence of the Gal4 Trans-Activator. Cell 52, 169–178.

Wu, Y., Reece, R.J., and Ptashne, M. (1996). Quantitation of putative activator-target affinities predicts transcriptional activating potentials. EMBO J 15, 3951–3963.

Zhu, G., Xia, Y., Nicholson, L.K., and Sze, K.H. (2000). Protein dynamics measurements by TROSY-based NMR experiments. J Magn Reson 143, 423–426.

