## Supplementary figures and images for "Mediator subunit Med15 dictates the conserved “fuzzy” binding mechanism of yeast transcription activators Gal4 and Gcn4"

### Supplemental Figures

#### Slide 1
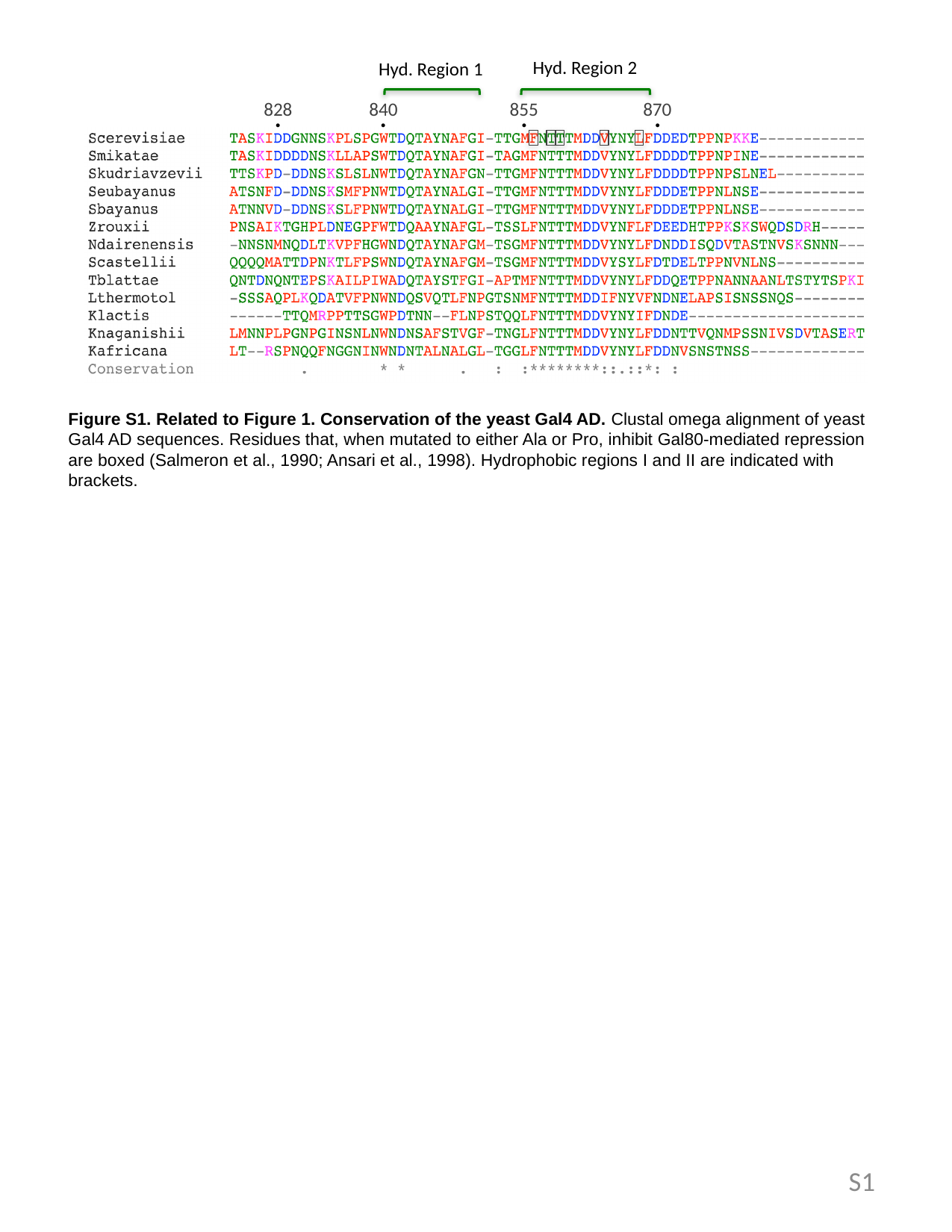

#### Slide 2
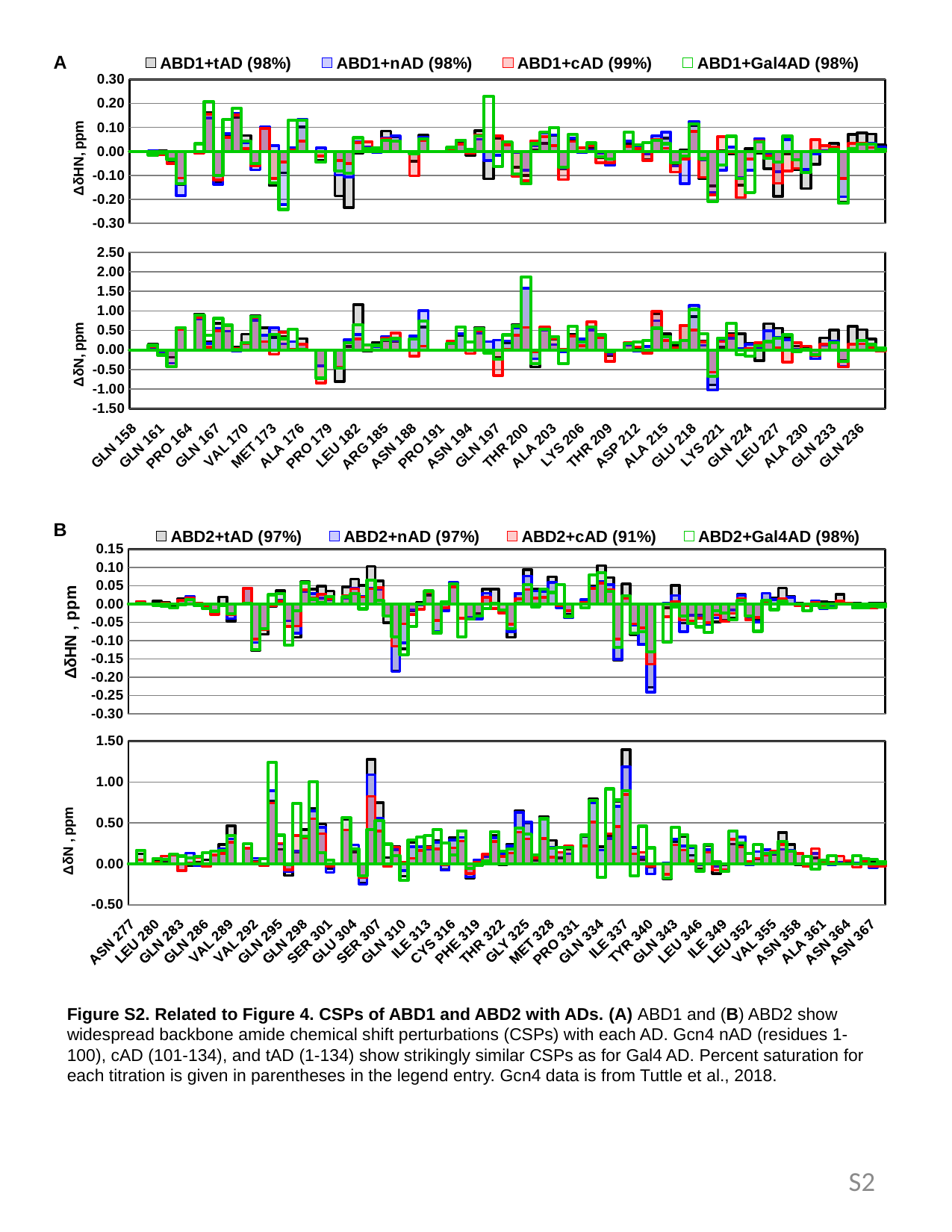

#### Slide 3
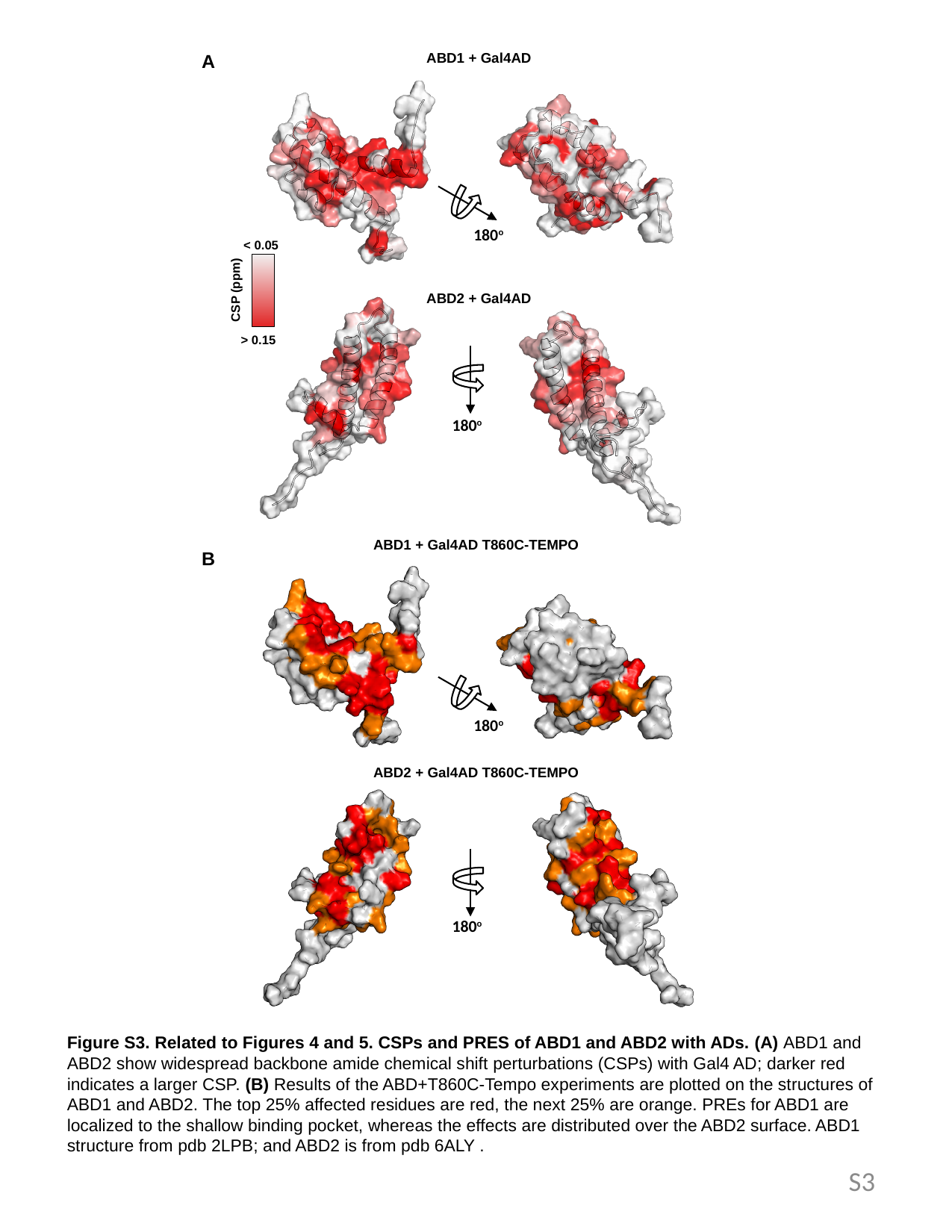
